# *Coxiella burnetii* infects osteoclasts and alters their differentiation and function in a type IV secretion system-dependent manner

**DOI:** 10.1101/2025.10.07.680913

**Authors:** Chaobo Lai, Md Nur A Alam Siddique, Faiza Asghar, Xudong Su, Jan Schulze-Luehrmann, Yewei Jia, Edith Alexandar Escarrega, Aline Bozec, Roland Lang, Anja Lührmann, Didier Soulat

## Abstract

Chronic Q fever is caused by persistent infection with the Gram-negative bacterium *Coxiella burnetii*. The mechanisms underlying this persistence remain elusive, but the presence of the bacteria in the bone marrow of *C. burnetii*-infected patients has been demonstrated. Therefore, we investigated the potential role of osteoclasts, the bone-resorbing cells, in harboring *C. burnetii* during the infection. The histological analysis of bones from a murine model of Q fever revealed the presence of *C. burnetii* inside osteoclasts. *In vitro* infection assays confirmed that osteoclasts can be infected with *C. burnetii* and supported bacterial replication in a type IVB secretion system (T4BSS)-dependent manner. Wild-type *C. burnetii* infection inhibited osteoclast differentiation and bone-resorbing activity, while the T4BSS mutant enhanced the differentiation and bone-degrading function of osteoclasts. Taken together, our findings identify osteoclasts as a potential host cell for *C. burnetii*, opening new perspectives on mechanisms that may underlie chronic Q fever. Additionally, infection-induced alterations in osteoclast function raise the possibility of alterations of the bone structure in affected patients.

## 2 Introduction

*Coxiella burnetii* is a Gram-negative, obligate intracellular bacterium and the causative agent of the disease Q fever (1). This zoonotic disease can be found worldwide, with the exception of New Zealand and Antarctica. It affects humans, as well as different animal species, like mammals, birds and ticks (2). Infected small ruminants bear a risk for the transmission of the disease to humans while showing a quite diverse picture of disease outcome, from asymptomatic to reproductive disorders and abortion (3). Infected ruminants shed the pathogen into the environment via feces, milk and mainly birthing products, such as amniotic fluid and placenta. The main route of human infection is via inhalation of contaminated aerosols and dust. While around 60% of infected humans are asymptomatic, acute disease manifestation spans from flu-like illness to pneumonia or hepatitis (2). Although most patients clear the infection, 20% of patients with a symptomatic Q fever infection will develop Q fever fatigue syndrome (QFS). This can last 5 to 10 years, and is characterized by severe fatigue, musculoskeletal pain, sleeping problems, impaired concentration and headache (4–6). So far, treatment options for QFS are limited and concentrate on cognitive behavioral therapy (7). Furthermore, 2 to 5% of all infected individuals develop chronic Q fever months or years after the primary infection. This chronic disease is characterized by endocarditis or vasculitis, mainly in patients with underlying valvulopathy (2). Treatment of chronic Q fever requires administration of doxycycline in combination with hydroxychloroquine for at least 18 months (8). This long treatment indicates that new therapeutic strategies have to be developed.

Tissue-resident alveolar macrophages of the lung are believed to be the first cells taking up the aerosolized bacteria (2). However, during the course of an infection the pathogen also infects other cell types, e.g., endothelial cells, fibroblasts, trophoblasts and epithelial cells (9). Uptake by phagocytic cells is accomplished by the α_v_β_3_ integrin and the complement receptor 3 (10, 11). Invasion into non-phagocytic cells is mediated by the *C. burnetii* outer membrane protein (Omp)A and the host cell protein CD44 (12, 13). In all infected cells, *C. burnetii* remains inside a phagolysosomal-like vacuole named the *C. burnetii*-containing vacuole (CCV). However, the maturation of the CCV diverges from the canonical phagosome maturation process by fusing early on with autophagosome and at later stages with secretory vesicles. In addition, *C. burnetii* delays the maturation of the CCV (14–16). Nevertheless, fully matured CCVs have phagolysosomal characteristics (17). The formation of such a mature phagolysosome would result in the destruction of most microbes and the activation of the immune system (18). In contrast, *C. burnetii* survives within this hostile environment and takes advantage of the acidic pH to activate one important virulence factor, the type IVB secretion system (T4BSS) (9, 19). The T4BSS is essential for its intracellular replication (20, 21). T4BSSs are multi-protein complexes used to inject bacterial effector proteins into the host cell cytosol to modulate host cell pathways in favor of the pathogen. For most *C. burnetii* T4BSS effectors the biochemical and molecular functions are unknown. Those studied to date modulate cellular processes such as autophagy, vesicular trafficking, gene expression, signaling or cell death (9, 22). For the latter, only anti-apoptotic (AnkG, CaeA and CaeB) and anti-pyroptotic (IcaA) effector proteins have been identified (23–28). These pro-survival functions are in line with the fact that inhibition of host cell death is essential for *C. burnetii* to complete its lengthy replication cycle, which takes ∼20 hours (2, 9).

The early stage of the infection is mostly asymptomatic (29), suggesting that *C. burnetii* persist unrecognized from the innate immune system within the human body, a fact that was already recognized since the 1940s (30). However, information about the site and regulation of *C. burnetii* persistence are still rare. Several reports indicated that *C. burnetii* antigen and/or DNA are present in the bone marrow of infected mice and humans months to years after infection (31–33). Indeed, the hypoxic environment of the bone marrow with an oxygen concentration ranging from 0.6% to 3% (34) corresponds to the conditions favoring *C. burnetii* persistence (35, 36). Therefore, we hypothesized that *C. burnetii* can persist in the bone marrow, as it has been demonstrated for several other human pathogens. Viruses such as HIV, bacteria like *Mycobacterium tuberculosis* and even parasites like *Plasmodium falciparum* or *Leishmania infantum* use the bone marrow niche to perpetuate their host infection (37–40).

Such infections of the bone marrow may alter the fine-tuned balance maintaining homeostasis of this organ. Indeed, bones are under continuous renewal to maintain their structure and functions. This process requires the antagonist activities of two cell types. While osteoblasts differentiating from mesenchymal precursor cells participate in bone formation by becoming osteocytes, osteoclasts of myeloid origin carry a bone resorbing function (41). Depending on the pathogen, a variety of bone marrow cell types have been identified as pathogen reservoirs. While macrophages represent the main cellular reservoir for intracellular pathogens, other cell types including stem cells populating the bone niche can play this role. Mesenchymal and hematopoietic stem cells (HSC) can both support silent infection with *M. tuberculosis* (38, 42). However, HSC have been more frequently reported as a cellular niche for various pathogens like virus with HIV (43) or parasite with *Leishmania* (44). Recent studies on this later pathogen illustrated the variety of consequences that infection can have on bone biology. During acute visceral leishmaniasis, the inflammatory response triggered by the infection dramatically alters the hematopoiesis taking place in the bone marrow leading to the expansion of HSC subsequently differentiating into myeloid cells permissive to the infection (45, 46). During the persistent phase of the disease, myeloid cells represent a safe niche for the parasite to persist in the bone marrow. In addition to macrophages, another phagocytic cell of the bone marrow has been identified as a safe niche for this intracellular parasite: the osteoclast (40). Like macrophages, these cells differentiate from the common myeloid progenitor before fusing into large multinucleated cells that are responsible for bone resorption. *C. burnetii*, like *Leishmania infantum*, is an intracellular pathogen particularly adapted to manipulate professional phagocytic cells. Therefore, we extended our hypothesis by suggesting that persistence of *C. burnetii* could be supported by osteoclasts in the bone marrow.

## 3 Materials and Methods

### 3.1 Bacteria strains

*C. burnetii* wild type (WT) corresponds to the Nine Mile Phase II strain (RSA439 clone 4) was kindly provided by Matteo Bonazzi (CNRS, Montpellier, France). The *C. burnetii* T4SS mutant used in this study has a genetic deletion of the dotA gene (Δ*dotA*) (47). The Δ*ankG* and Δ*caeB* strain were previously established and described in (47) and (26) respectively. Both bacterial strains were grown in acidified citrate cysteine medium-2 adjusted to pH 4.75 (ACCM-2, Sunrise Science Products) at 37°C with 5% CO_2_ and 2.5% O_2_ (48). The bacteria were inoculated at an OD_600_ of 0.01 and growth of the culture was monitored by spectrometry until it reached 0.2 - 0.3, which is usually around day 4-5 after inoculation. Heat killed WT *C. burnetii* were prepared from this mature culture. Bacteria were first washed with sterile PBS. Aliquots of 20 µL were prepared in 200 µL PCR tube with a bacterial suspension at 5×10^8^ mL ^-1^. Then, the bacteria were heated for 2 min at 65°C, incubated on ice for 5 min and finally incubated at 37°C for 5 min and stored on ice.

### 3.2 Animals and infection

All mice utilized in this study were at least 6 weeks of age. C57BL/6 wild-type mice were obtained from Charles River Breeding Laboratories (Sulzfeld, Germany) or bred at the Präklinische Experimentelle Tierzentrum of the University Hospital Erlangen (PETZ). *Myd88*^−/−^ mice (Myd88 tm1Aki) were provided by Dr. S. Akira (University of Osaka, Japan). *Rag2*-/-*γ-Chain*-/-mice were kindly provided by Dr. Ulrike Schleicher (Erlangen). All mouse experiments were approved by the regional government (Regierung von Unterfranken, animal protocols 54-2532.1-44/13 and 55.2.2– 2532.2-854-14). Mice were bred at the PETZ of the University Hospital Erlangen and transferred to a biosafety level 2 animal room at least 1 week before infection. Both female and male mice were used in the infection experiments. Groups of mice were infected with NMII intraperitoneally (1×10^7^ CFU/200 μL PBS/mouse) as described before (49). The physical condition of the mice was monitored regularly, including measuring the weight of the animals. At the indicated time points, mice were sacrificed by cervical dislocation and tibia bone were collected.

### 3.3 Immunofluorescence in bone sections

Tibia bones were fixed in 4% paraformaldehyde (PFA, Alfa Aesar) overnight and decalcified for 7 days in 14% EDTA. Bones were then incubated for 12 h in 30% sucrose in PBS before being embedded in Tissue-Tek® O.C.T compound (Science Services) for cryo-sectioning (6 µm). For the immunofluorescence staining of *C. burnetii* and CD68 we proceeded as follows. Bone sections were permeabilized with 0.1% Triton X-100 in PBS for 20 min at room temperature (RT). Sections were then washed three times with PBS and incubated in 10% goat serum in PBS for 1h at RT. After three washing steps of 5 min with PBS, sections were incubated with a primary Ab diluted in 0.5% goat serum in PBS overnight at 4°C, followed by another three washing steps and an incubation with a suitable secondary Ab diluted in 0.5% goat serum in PBS for 1 h at RT in the dark. After three final washings, sections were mounted with Molecular Probes™ ProLong™ Diamond Antifade Mountant (Life Technologies), containing DAPI to stain DNA and cured overnight. For the immunofluorescence staining of *C. burnetii* and TRAP we proceeded as follows. Bone sections were permeabilized with 0.2% Tween 20 in PBS for 20 min at RT. Sections were stained with the primary and secondary antibodies as previously described. Before mounting, the staining for TRAP was realized with the ELF-97 Endogenous Phosphatase Detection Kit following manufacturer’s instructions (diluted 1:20, E6601, Invitrogen). Finally, sections were mounted with the mounting medium provided in the kit and cured overnight. Polyclonal rabbit anti-*C. burnetii* antiserum (1:2000) (50) and rat anti-CD68 (diluted 1:200; GTX43914, Genetex/Biozol), were used as primary antibodies. Goat anti-rabbit Alexa488 antibody (1:600; Jackson immune research, 111-545-045), goat anti-rabbit Alexa-594 (1:600, Jackson immune research, 111-585-045) and goat anti-rat-IgG (H + L) Dylight550 (diluted 1:200; ab96888, Abcam) were used as secondary antibodies. The immunofluorescence images were acquired with a confocal laser scanning fluorescence microscope (LSM700; Zeiss).

### 3.4 Quantification of *C. burnetii* burden in bone marrow via qPCR

Bone marrow was collected in 800 µL PBS from mice bone and weighted. Bone marrow cells were ruptured with 2 mm iron beads in an Omni Bead Ruptor 24 (Omni International) by 5 rounds of shaking at 2.9 m/s for 10 s interval by 5 s pause. 150 µL of the homogenate was centrifuged (20,000 ×*g*, 2 min, RT). 300 µL Lysis buffer (0.1 M EDTA, 0.1 M NaCl, 1% SDS, 0.05 M Tris/HCl (pH 8)) plus proteinase K (6 U/mL) was added to the pellet and incubated at 55°C and 900 rpm overnight. 300 µL isopropanol was added and mixed by inversion. The solution was centrifuged (20,000 ×*g*, 30 min, 10°C) to precipitate the DNA. This was followed by 2 washing steps with 70 % EtOH (20,000 ×*g*, 15 min, RT). The pellet was air-dried and dissolved in a.d. and incubated overnight at 4°C. The genomic DNA was used to defined the ratio of *C. burnetii* genomic copies to BMM genomic copies as bacterial load per cell (51). *C. burnetii* genomic copies were quantified with specific primers for the insertion sequence IS1111 (Table. 1). Host cell genomic copies were quantified from the same sample using a primer set specific for *Alb*, the murine albumin gene (Table. 1). Standard curves for murine genome was generated by serial dilution of DNA from mouse spleen cells. To quantitate *C. burnetii* genome equivalents, a standard curve using titrated DNA prepared from a defined number of *in vitro* cultured NMII was prepared.

### 3.5 Quantification of *C. burnetii* burden in bone marrow via CFU

Suspension of bone marrow cells were ruptured as described above. The homogenates were diluted 1 to 10 in series up to a dilution factor of 10^-5^. A volume of 5 µL of each dilution was dropped in triplicate on solid medium (ACCM-2, 0.3% agar). Plates were incubated at 37°C within a controlled atmosphere containing 2.5% O_2_ and 5% CO_2_. After 7 to 10 days, colonies were counted for each dilution to calculate the number of CFU per gram of tissue.

### 3.6 Murine bone marrow macrophage differentiation

Bone marrow macrophages (BMMs) were generated as previously described (52) from femur of C57BL/6N mice. For infection, BMM were seeded in RPMI-1640 (Gibco), supplemented with 10 mM HEPES, 50 µM 2-mercaptoethanol and 10% (v/v) heat-inactivated FCS in 24 well plates (4.0×10^5^ cells/well). BMM were infected with *C. burnetii* at a MOI 10 for 6 h before the bacteria were washed with warm PBS twice.

### 3.7 Murine osteoclast differentiation

Total bone marrow cells from wild-type (WT) mice were extracted by flushing the femur and tibia. The cells were then plated overnight in a 10 cm cell culture plate containing 10 mL of osteoclast medium (αMEM (Gibco), GlutaMAX (Gibco), 10% FCS, 1% penicillin/streptomycin (Gibco)) supplemented with 5 ng/mL M-CSF (PeproTech). The non-adherent cells were collected the next day and cultured in osteoclast medium with 20 ng/mL M-CSF and 20 ng/mL RANKL (PeproTech) in 96-well plates (for TRAP staining; 200 μL/well) or 24-well plates (for RNA preparation; 1 mL/well) at a concentration of 1×10^6^ cells/mL. The medium was renewed every 2 days. The cultures were maintained at 37°C under humidified atmosphere with 5% CO_2_.

Early and late differentiated osteoclasts (day 3 and day 5) were washed with PBS and fixed using 4% paraformaldehyde (PFA, Alfa Aesar).

### 3.8 Immunofluorescence in osteoclast

Osteoclasts, which were previously seeded on a glass coverslip in a 24-well plate, were infected with *C. burnetii* on day 2 of differentiation (6 h prior the 1st change of medium) at a multiplicity of infection (MOI) of 10. On 24 hpi and 72 hpi, the cells were washed with PBS and fixed with 300 µL of 4% PFA. The cells were first permeabilized with 0.05% Triton X-100 in H_2_O and then blocked with 5% goat serum in PBS. The cells were then stained for *Coxiella* using a rabbit anti-*Coxiella* antiserum (1:5,000) (50) in 0.5% of goat serum in PBS as the primary antibody. Goat anti-rabbit Alexa488 antibody (1:600 Jackson immune research 111-545-045) in 0.5% goat serum in PBS was used as a secondary antibody. In parallel to the secondary antibody, Actin was stained using Phalloidin-Alexa647 (1:200, Thermo Scientific). DNA was stained using DAPI incorporated in the mounting medium (ProLong™ Gold Antifade DAPI Mountant, molecular probes). Images were captured using an LSM700 confocal microscope. The bacterial burden was quantified by analyzing the images of infected osteoclasts or non-osteoclasts on stained coverslips using the Integrate density/Area “IntDen/Area” parameter in ImageJ, which represents the mean fluorescence intensity of bacteria in the Alexa488 channel divided by the area of the cells. Cells nuclei number (DAPI channel) were counted using imageJ.

### 3.9 TRAP staining

Histochemical TRAP staining (Sigma Aldrich) of osteoclast cultures in 96-well plates was performed at 3 and 5 days of differentiation following the manufacturer’s instruction. Briefly, the cells were washed with PBS before being fixed with 4% PFA for 2 minutes at 37°C. After two washes with warmed PBS, the cells were stained with 100 μL of the TRAP staining solution per 96-well plate and incubated for 10 minutes at 37°C. Following the staining, the cells were washed with PBS. Images of the 96-well plates were captured with a light microscope (Keyence). The number of osteoclast (TRAP+ and >3 nuclei/cell), their number of nuclei per cell, their diameter and the total number of cells were quantified using ImageJ.

### 3.10 TUNEL assay

DNA strand breaks were detected by incubating cells with TUNEL reaction mixture containing the terminal deoxynucleotidyl transferase and fluorescently labelled nucleotides using the manufacturer’s protocol (Roche, 12156792910). Osteoclasts and *C. burnetii* were stained following the above-described protocol (Immunofluorescence in Osteoclast). The number of TUNEL-positive cells was analyzed using an LSM700 confocal microscope.

### 3.11 Resorption assay

To perform the resorption assay, calcium phosphate (CaP) coated cell culture plates were prepared. Sterile solution of 0.12 M Na_2_HPO_4_ and 0.2 M CaCl_2_ at pH7.4 were pre-incubated overnight at 37°C to maximize solubilization. Next day, solutions were mixed in a 1:1 ratio and washed three times with sterile water after centrifugation for 10 min at 360 ×*g*. The pellet was then resuspended in sterile water with a 10 times dilution factor. 200 µL of the suspension was added to each well of a 96-well cell culture plate. The coated plates were then air-dried in a closed sterile hood at RT for 3-5 days until fully dried. The osteoclast culture could then be initiated following the previously described protocol. Cells were infected on day 2 of differentiation (6 h prior the medium change). After 7 days of differentiation, the osteoclasts were lysed using deionized water and incubated with 5% of sodium hypochlorite (MilliporeSigma) for 5 minutes. This treatment removed the cells and revealed the resorbed area. Images were acquired with a light microscope (Keyence). The quantification of the resorbed area was performed by calculating the averaged percentage of the resorbed area using ImageJ.

### 3.12 RNA extraction and quantitative RT-PCR

Total RNA from osteoclasts or BMM were isolated using TRIzol (Invitrogen) or RNA-Solv® Reagent (VWR) according to the manufacturer’s instructions. mRNA samples were reverse transcribed into cDNA using an oligo(dT) primer and a reverse transcriptase (Thermo Scientific). Quantitative real-time PCR (qPCR) was performed using SYBR Select Master Mix (Lifetech). Samples were analyzed in triplicate. mRNA expression was reported as relative expression (2^-ΔCt^) or fold induction (2^-ΔΔCt^) and β-actin were used to normalize the RNA content of samples. Primer sequences are listed in table 1.

**Table 1.**
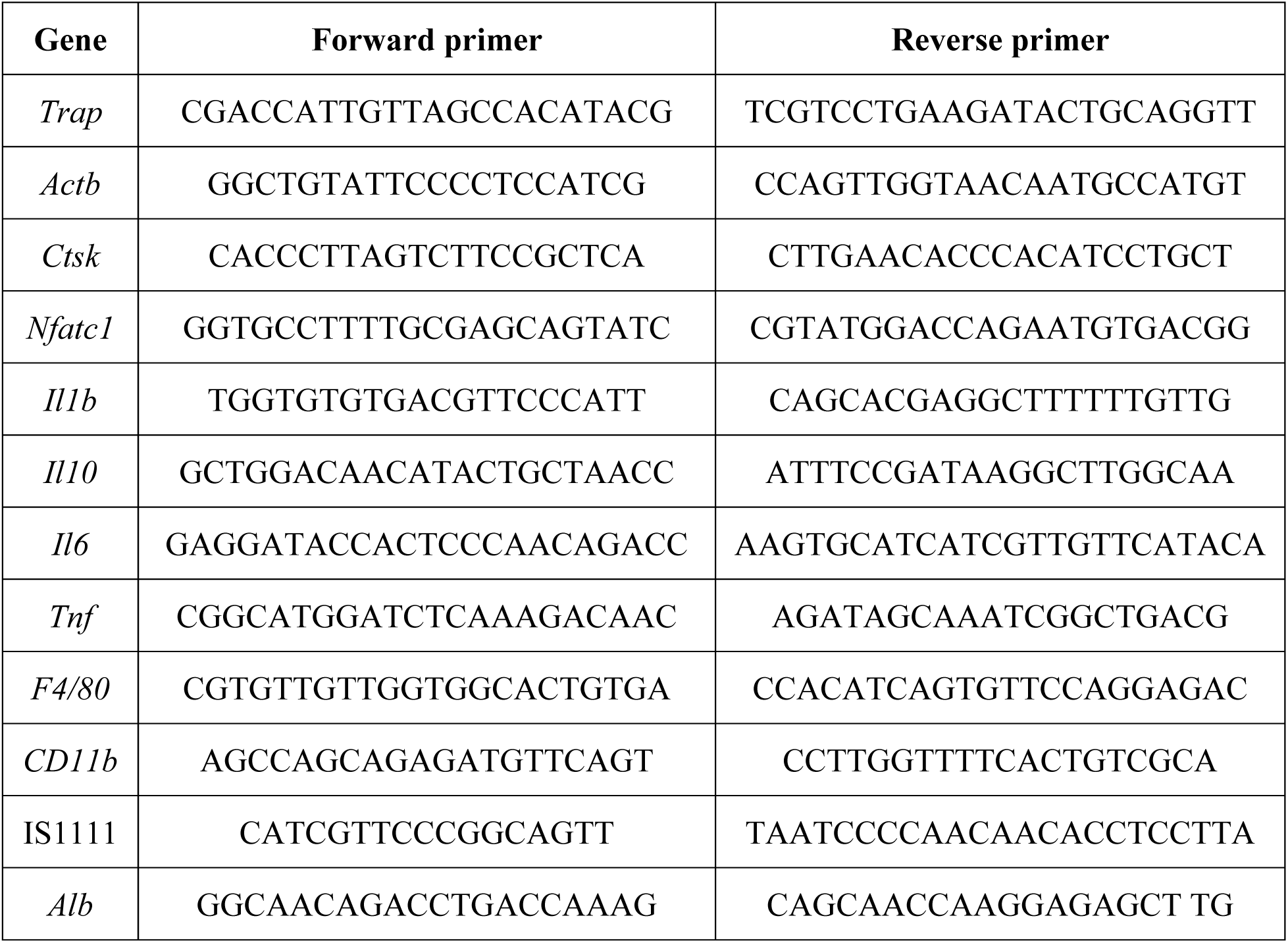
Sequence of the primers used for qPCR.

### 3.13 Enzyme-linked immunosorbent assay (ELISA)

Osteoclast supernatants were used for the measurement of TNF, IL-6, IL-1B and IL-10 protein levels. The following ELISA kits were used according to the manufacturer’s instructions: the BD OptEIA Mouse TNF ELISA Set (BD Bioscience; Cat# 555268), BD OptEIA Mouse IL-6 ELISA Set (BD Bioscience; Cat# 555240), the Mouse IL-1 beta/IL-1F2 DuoSet ELISA (R&D; cat# DY401-05), the BD OptEIA Mouse IL-10 ELISA Set (BD Bioscience; Cat# 555252).

### 3.14 Statistics

All data are presented as mean ± SD. The statistical significance was determined either by unpaired student’s t test (normal distribution) or Mann Whitney test for comparing two data sets using GraphPad Prism software 9.0. The p-value of the effect of each independent variable and their interaction is indicated on the figure when relevant.

## 4 Results

### 4.1 *C. burnetii* infection results in their capture by osteoclasts in the bone marrow

Wild type (WT) laboratory mouse strain proved to be resistant to *C. burnetii* infection via the natural route of infection as they quickly clear the bacterial infection. In humans, polymorphism of *MyD88*, an adaptor shared by most TLRs and the IL1 receptor, has been linked to Q fever susceptibility (53). Similarly, the genetic deletion of *Myd88* proved to be sufficient to establish a murine model permissive to *C. burnetii* infection (49). Taking advantage of this model, we analyzed the spreading of *C. burnetii* Nine Mile phase II (WT) into the liver, spleen and bone marrow of *Myd88* deficient and WT mice five days after intra-peritoneal infection, using non-infected mice as control. The number of living *C. burnetii* per gram of all tested tissue (Figure 1A) was higher in MyD88 deficient mice compared to infected control mice, confirming the adequacy of the model (49). Cryosection of tibial bone were stained for *C. burnetii* and CD68, also named macrosialin, which is a surface protein expressed by macrophages and osteoclasts in the bone marrow. Confocal laser scanning fluorescence microscopy (CLSFM) analysis of the bone of *Myd88* deficient mice revealed the presence of *C. burnetii* inside multinucleated cells lining bone trabecula and expressing CD68, three characteristics defining osteoclasts (Figure 1B). To confirm that the detection of these events was reliable, we also infected the highly susceptible mouse strain *Rag2*-/-*γ-Chain*-/-for 14 days. Indeed, the quantification of the immunohistofluorescence staining of the bone confirmed the reliability of the *Coxiella* detection as we observed a substantial increase of infected CD68+ cells in the bone marrow of these immunocompromised mice compared to control mice (Figure 1C). To substantiate the putative infection of osteoclast with *Coxiella*, additional bone sections from infected *Myd88* deficient mice were stained simultaneously for *Coxiella* and for the expression of a marker of osteoclast terminal differentiation, the tartrate-resistant acid phosphatase (TRAP) and compared to sections of infected and uninfected control mice (Figure 1D). This staining revealed that the proportion of infected osteoclasts among osteoclast (Figure 1E) or among infected cells (Figure 1F), including TRAP negative macrophages and other phagocytes, were comparable between control and *Myd88* deficient mice. These data suggest that cells of the bone marrow and specifically osteoclasts can be infected by *C. burnetii in vivo*.

**Figure 1.**
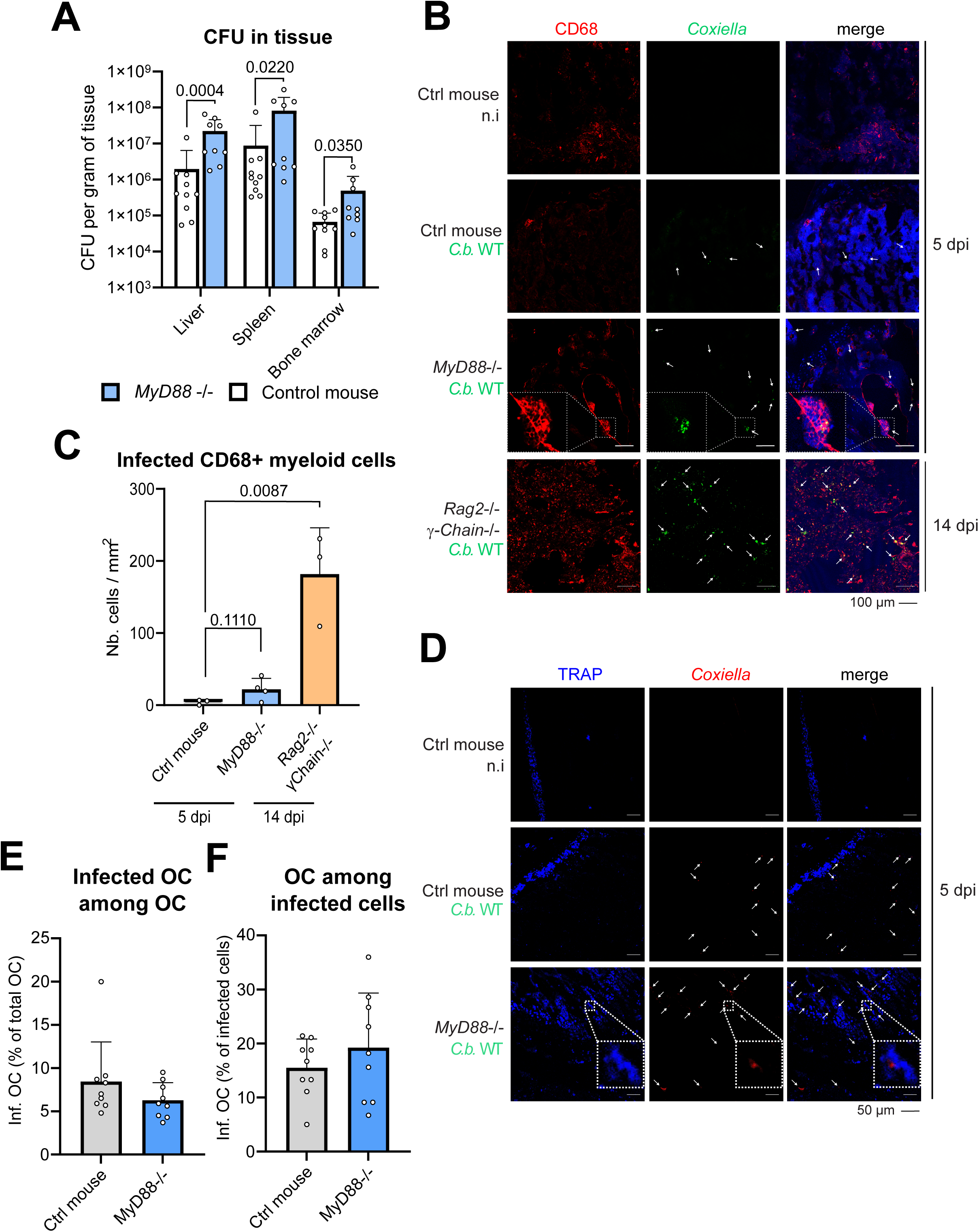
***C. burnetii* can infect bone marrow myeloid cells including osteoclasts *in vivo*. (A)** *C. burnetii* was injected intraperitoneally (i.p.) in control (ctrl) or *Myd88* knock out mice (1.0 × 10^7^ b/mice). The number of living *C. burnetii* in liver, spleen and bone marrow (CFU) of these mice was measured by limiting dilution (n = 10 for ctrl; n=9 for knock out mice). (B) Control (ctrl), *Myd88*-/-or *Rag2*-/-*γ-Chain*-/-mice were similarly infected i.p. for 5 or 14 days. The localization of *C. burnetii* in myeloid cells of the BM was analyzed in the femurs of these mice (n = 6 for ctrl and *Myd88*-/-mice; n=3 for *Rag2*-/-*γ-Chain*-/-mice). Bone tissues were cryosectioned and stained for *Coxiella* (green-Alexa488), CD68 (expressed by osteoclasts and macrophages; red-Alexa555), and DNA (blue-DAPI). Sections were imaged by CLSFM. Foci of infection are marked with white arrows. Inserts of infected CD68 expressing cells lining a trabecula are depicted within dashed line. Scale bar represents 100 µm. (C) Number of infected CD68+ myeloid cells per mm^2^ of bone marrow section was quantified (n=3 to 4 mice for each condition). (D) The co-localization of *C. burnetii* with osteoclast was analyzed in the femurs of ctrl and *Myd88*-/-infected mice. Bone tissues were cryosectioned and stained for *Coxiella* (green-Alexa594) and TRAP (expressed by osteoclasts only; blue-ELF97). Sections were imaged by CLSFM. Foci of infection are marked with white arrows. Inserts of infected osteoclast are depicted within dashed line. Scale bar represents 50 µm. (E) Percentage of infected osteoclast among all osteoclast was quantified (n=3 mice for each condition). (F) Percentage of osteoclasts among all infected cells of the BM was quantified (n=3 mice for each condition). Data are shown as mean ± SD.

### 4.2 *C. burnetii* infect osteoclasts *in vitro*

To investigate whether *C. burnetii* influences osteoclast differentiation and function, we established a *C. burnetii in vitro* infection model of osteoclasts (Figure 2A). Bone marrow progenitor cells were initially cultured in presence of macrophage colony-stimulating factor (M-CSF) to amplify the pool of myeloid progenitors and to discard other cell types. Next, the progenitor cells were differentiated for 2 days in presence of M-CSF and receptor activator of nuclear factor kappa-Β ligand (RANKL) before being infected with *C. burnetii* (WT). At this stage, osteoclasts are progressing toward their multinucleated and matured form that they would reach by day 3 to day 4 of differentiation. We decided to not infect fully matured osteoclasts because of their limited viability after 5 days of differentiation, particularly for non-infected cells. After 24 h and 72 h of infection, cells were stained for *C. burnetii* and Actin to visualize the cell cortex and therefore segregate the cells from each other. At 24 h post-infection, CLSFM analysis confirmed that osteoclasts could phagocytose *C. burnetii* (Figure 2B, left panel). In addition to multinucleated osteoclasts, these *in vitro* cultures contained non-fused myeloid cells that resemble macrophages due to the continuous M-CSF stimulation. These phagocytic cells also harbored substantial quantity of bacteria. Calculating the ratio between the staining intensity of phagocytosed *C. burnetii* and the cell surface, we estimated that these cells contained 8 times more bacteria than osteoclasts at 24 h post-infection (Figure 2B and D). As *C. burnetii* might replicate intracellularly only once within the first 24 h of infection (2), we concluded that osteoclasts phagocytose the bacteria, but to a lesser extent than macrophages.

**Figure 2.**
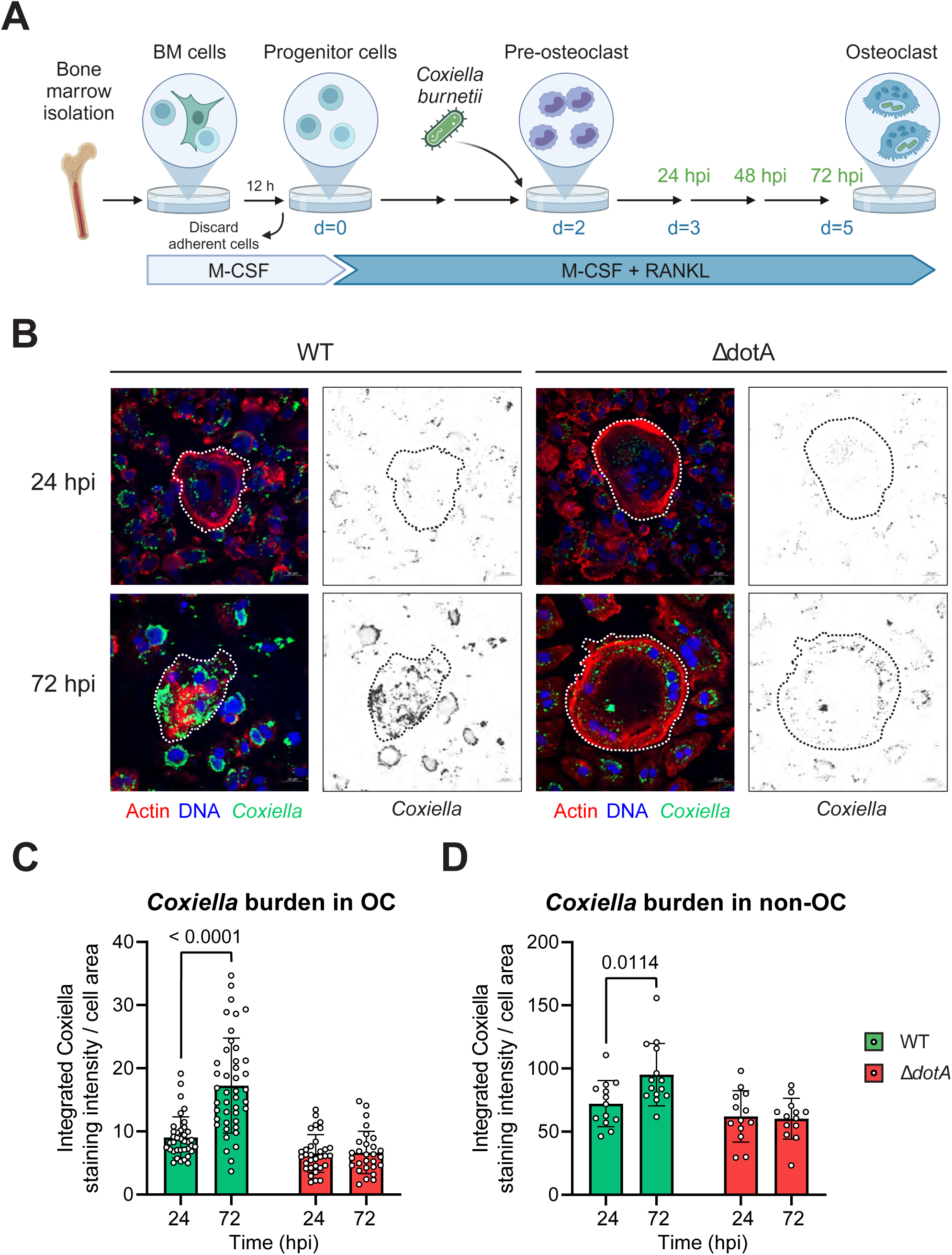
***C. burnetii* needs dotA to sustain its growth in myeloid cells including osteoclasts. (A)** Osteoclasts were differentiated from bone marrow progenitor cells in presence of M-CSF and RANKL. Cells were then infected after 2 days of differentiation with *C. burnetii* WT or Δ*dotA* at MOI 10. Uninfected osteclasts (n.i.) were used as control. **(B)** Osteoclasts differentiated from bone marrow progenitor cells (Fig. 1A) were infected after 2 days of differentiation with *C. burnetii* WT or Δ*dotA* at MOI 10. After 24 or 72 hpi, cells were stained for *Coxiella* (green - Alexa488), actin (red – Alexa647) and DNA (blue - DAPI) and imaged by fluorescent CLSFM (n=3). The bacterial burden was quantified using imageJ to calculate a ratio of the measured intensity of the staining emitted by *C. burnetii* within each cell divided by the measured surface of this cell. The scale bar represents 20 µm. The bacterial burden was analyzed for the osteoclasts **(C)** and for the other myeloid cells **(D)**. Data are shown as mean ± SD. Circle represent individual cells.

### 4.3 *C. burnetii* replication in osteoclasts *in vitro* requires the T4BSS

Focusing on osteoclasts, it appears that *C. burnetii* can thrive in these cells as we detected a significant increase of the bacteria at 72 h post-infection. Within this 48 h interval, the number of *C. burnetii* doubled within osteoclasts (Figure 2B and C). Interestingly, the macrophage, which initially phagocytosed more bacteria, showed a significant, but substantially smaller increase of intracellular bacteria (Figure 2B and D). Therefore, osteoclasts appear to be more permissive for *C. burnetii* replication and / or survival than mononuclear phagocytic myeloid cells.

Next, we investigated whether *C. burnetii* also relied on its T4BSS to infect and replicate inside osteoclast, like it does for macrophage (54). Thus, we infected osteoclasts with a T4BSS-defective mutant, Δ*dotA C. burnetii*, and with the wild type strain. For the earliest time point, both strains could infect osteoclasts similarly well (Figure 2B and C). Likewise, other non-phagocytic cells took up indifferently both strains (Figure 2B and D). However, the *C. burnetii* Δ*dotA* failed to replicate inside osteoclast and other phagocytic cells (Figure 2C and D). Therefore, we concluded that intracellular replication of *C. burnetii* inside osteoclast depends on a functional T4BSS.

### 4.4 Infection with *C. burnetii* inhibits osteoclastogenesis *in vitro*

Next, we studied the consequence of the *C. burnetii*-infection on osteoclasts. First, osteoclast differentiation was quantified in uninfected and infected cultures by staining for the expression of TRAP. The expression of this enzyme and the presence of 3 or more nuclei per cell were the criteria used to define mature osteoclasts (Figure 3A). In absence of infection, a relatively high number of cells fused to form osteoclasts after 3 days of differentiation, the time point corresponding to 24 hpi for the infected samples. These non-infected cells were initially relatively small as they contained five nuclei within a diameter of 73 µm (Figure 3B-D). Over the next 2 days of differentiation, these osteoclasts grew in size attaining 14 nuclei per cell within a diameter of 268 µm. Interestingly, the number of osteoclasts observed for this condition remained unchanged probably because of a balance between two fusion processes having opposite influence on the absolute number of osteoclasts. While, fusion between single nucleated cells increased the osteoclast number, fusions between osteoclasts themselves reduced the osteoclast number.

**Figure 3.**
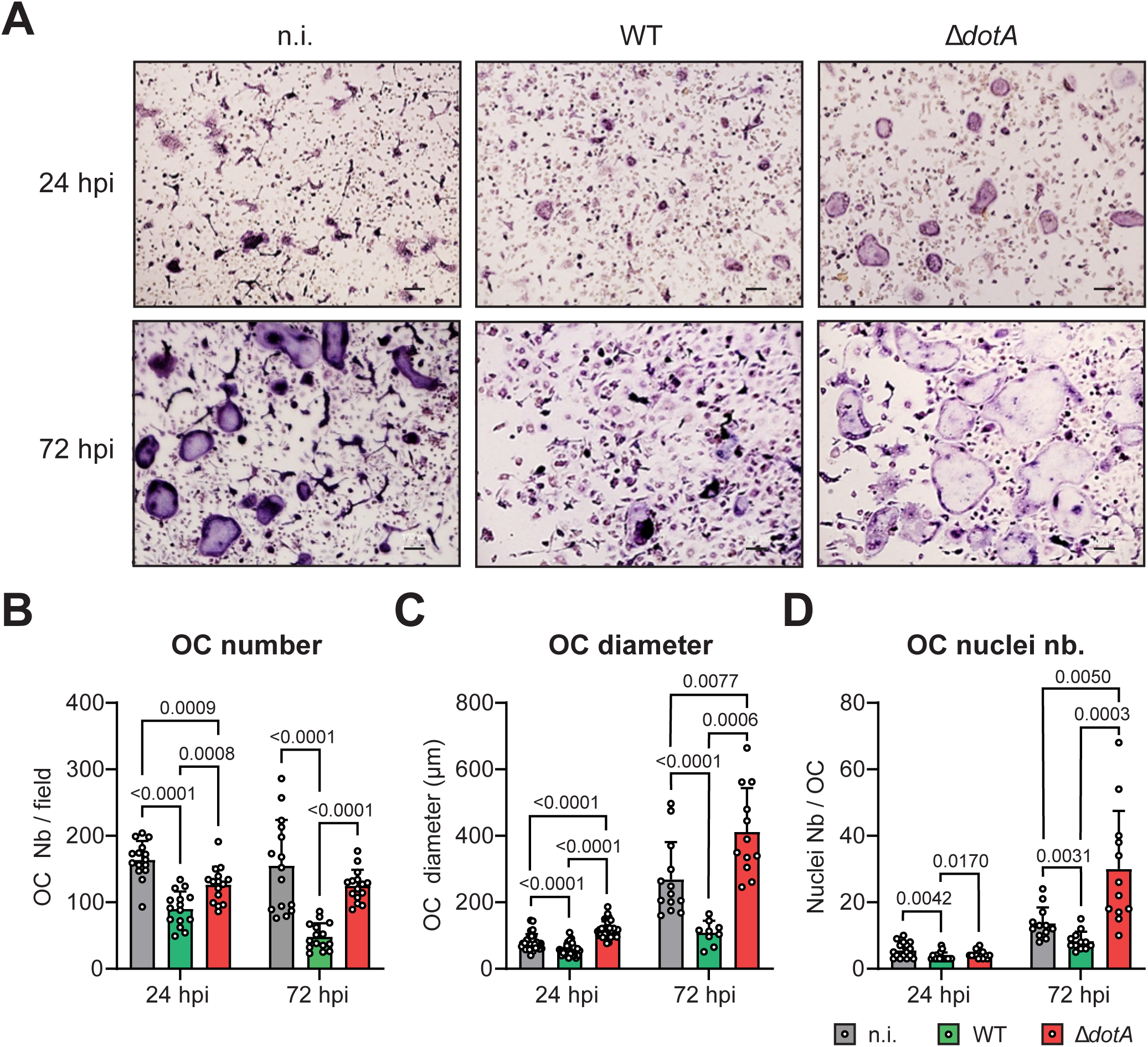
***C. burnetii* infection inhibits osteoclast differentiation in a DotA dependent manner. (A)** Osteoclast cultures were infected with *C. burnetii* WT or Δ*dotA* after 2 days of RANKL stimulation. Osteoclast differentiation was quantified 24 hpi and 72 hpi by staining for TRAP. Cells were analyzed by light microscopy. Scale bar represents 100 μm. **(B)** Osteoclasts (TRAP+ cells with ≥3 nuclei) were counted with imageJ software (n=3). **(C)** Similar approaches were used to measure the osteoclast diameter and **(D)** the number of nucleus per field as a proxy for cell survival. Error bars represent the standard deviation. Data are shown as mean ± SD. Circle represent individual microscopical fields of view.

Infection of osteoclasts with WT *C. burnetii* drastically inhibited their differentiation. Already at 24 h post-infection, which correspond to 3 days of differentiation, the osteoclast density, the number of nuclei and the size (55 µm) was significantly reduced compared to non-infected cells (Figure 3B-D). This general inhibition was accentuated at 72 h post-infection, when three times less osteoclasts were counted. In addition, these osteoclasts harbored half the number of nuclei (8 nuclei per cell) and were 2.5 times smaller than the non-infected cells. Strikingly, infection of osteoclasts with a DotA-deficient strain of *C. burnetii* had opposite consequences. While being only slightly reduced in number by 23% at 24 h post-infection, osteoclasts infected with *C. burnetii* Δ*dotA* had similar number of nuclei (4 nuclei per cell) and harbored even a higher diameter (114 µm) than non-infected cells (Figure 3B-D). This phenotype was enhanced at 72 h post-infection. After 3 days of infection with the mutant strain, osteoclasts harbored an excessively high number of nuclei (30 nuclei per cell) and developed in significantly larger cells (412 µm) than the non-infected cells or the cells infected with the WT bacteria. Surprisingly, this increased differentiation did not result in a higher number of osteoclasts when compared to non-infected cells. Nonetheless, these results illustrated the negative impact that *C. burnetii* infection has on osteoclast differentiation *in vitro* and the importance of the T4BSS in this process.

While infection inhibits undergoing osteoclastogenesis, we hypothesized that an earlier infection of monocytic precursors could have a more drastic effect on this process. Therefore, precursor cells grown only with M-CSF were infected with *C. burnetii* WT or Δ*dotA* for 6 h before being stimulated with RANKL to initiate osteoclastogenesis. Osteoclast differentiation was quantified by TRAP staining after 3 (72 hpi) and 5 days (120 hpi) of differentiation like for the previous experiment (Figure 4A). As illustrated in figure 4B, osteoclastogenesis was totally abrogated by infection with the WT or the mutant strain of *C. burnetii*. While the average size (Figure 4C) and number of nuclei per osteoclast (Figure 4D) appears to be similar in infected and non-infected cells, it is important to keep in mind that the values of the infected samples correspond to a very limited number of cells. Therefore, infection of monocytic precursor cells by *C. burnetii* appears to largely inhibit osteoclastogenesis without altering the fate for the rare cells that could enter the process of differentiation.

**Figure 4.**
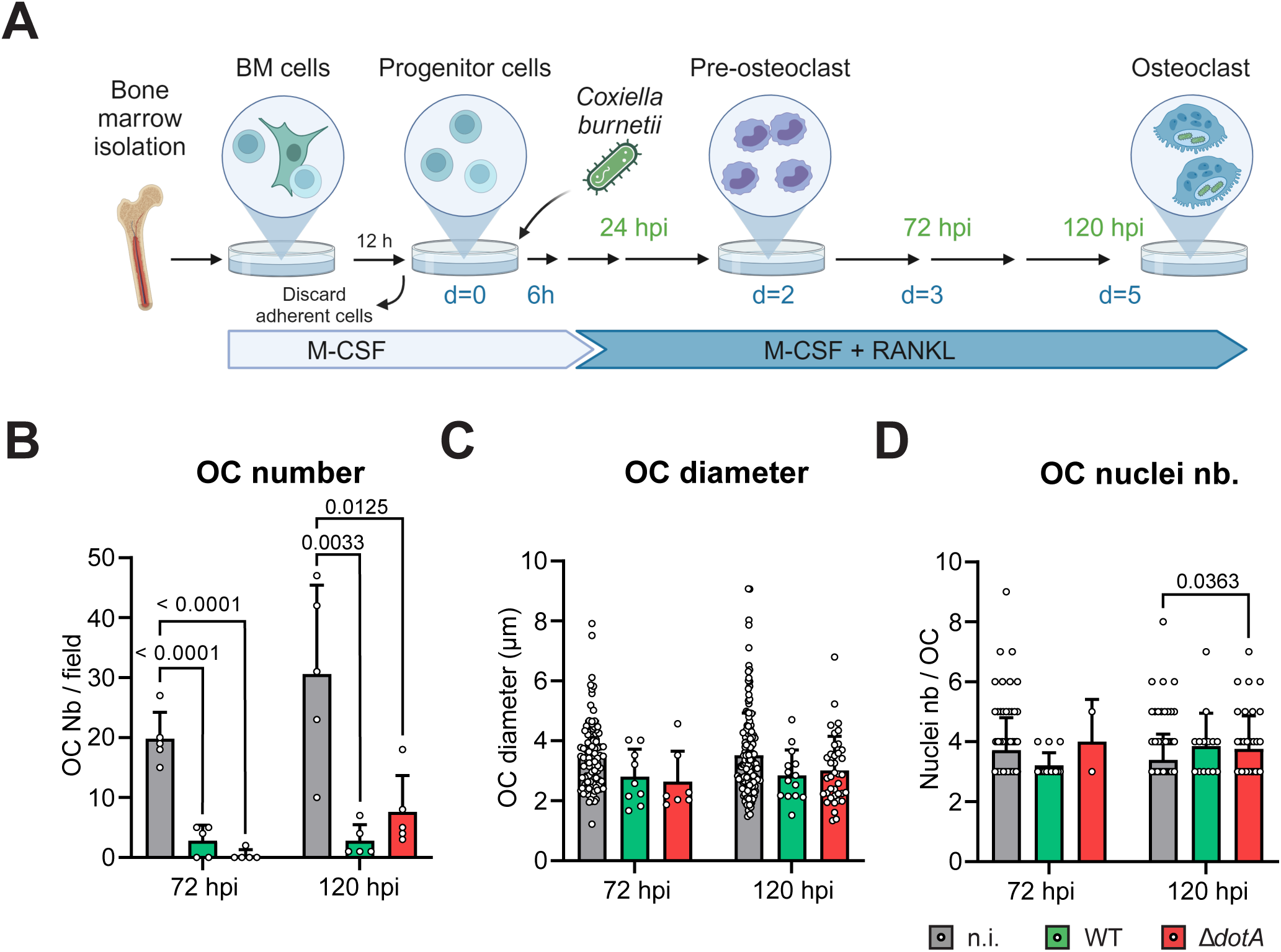
***C. burnetii* infection of myeloid progenitor cells inhibits osteoclast differentiation. (A)** Osteoclast progenitors were infected with *C. burnetii* WT or Δ*dotA* before being stimulated with RANKL (6 hpi). Osteoclast differentiation was analyzed 72 hpi and 120 hpi (matching time point with the previous experiment presented in Figure 3A) by staining for TRAP. **(B)** Osteoclasts number, **(C)** diameter and **(D)** nuclei number were quantified (n=2). Error bars represent the standard deviation. Data are shown as mean ± SD. Circle represent individual microscopical fields of view.

The differentiation of osteoclasts is a biological process governed by defined transcription factors such as Nuclear factor of activated T-cells, cytoplasmic 1 (NFATc1) that controls the expression of enzymes critical for the bone resorption function of osteoclasts such as the tartrate-resistant acid phosphatase type 5 (TRAP) and Cathepsin K. Knowing that *C. burnetii* infection altered the development and fusion of osteoclasts, we investigated if the infection of differentiating osteoclasts would also influence the expression of these functional markers of differentiation. While the expression of *Nfatc1* gene peaked after 4 days of differentiation in non-infected osteoclasts (48 h post-infection), it peaked earlier in cells infected with WT *C. burnetii*, namely at 6 h post-infection (Figure 5A). Moreover, *Nfatc1* expression was significantly lower in these cells from 24 h post-infection onward when compared to non-infected cells. Likewise, the expression of the phosphatase gene *Trap* and the protease gene *Ctsk* were similarly reduced in osteoclasts infected with WT bacteria compared to non-infected cells from 24 h post-infection onward (Figure 5B and C). This general reduction of expression of osteoclast marker genes after infection with WT *C. burnetii* was not observed in cells infected with the Δ*dotA* mutant. On the contrary, these cells expressed significantly more *Nfatc1* than non-infected cells at 24 h post-infection. This trend, despite being not significant, was also observed for the expression of *Trap* and *CtsK* at 24 h post-infection. Nonetheless, the Δ*dotA*–induced early and excessive expression of differentiation markers was followed by a strong reduction of the transcription of *Nfatc1* and *Trap* from 48 h post-infection onward (Figure 5A and B), while *Ctsk* expression returned to values of non-infected cells (Figure 5C). As the osteoclast culture is by essence a mix of cells comprising osteoclast and differentiating monocytic precursor cells / macrophages, we aimed to quantify their specific differentiation and infection status. Although a FACS-based approach has been considered, this technique is not suitable for our experimental setup due to the infection-induced fragility of these multinucleated cells. Therefore, we addressed the expression of macrophage markers in this culture to evaluate the degree of differentiation. Expression of *Itgam* and *Adgre1*, coding for CD11b and F4/80 respectively, were quantified in osteoclast infected with *C. burnetii* WT and Δ*dotA* 24 hpi and compared to bone marrow derived macrophages treated similarly. Osteoclast cultures expressed significantly less of both genes than macrophages when cells were not infected (Figure 5D and E). This pattern tends to stay true for CD11b when cells were infected (Figure 5D). Interestingly, expression of *Adgre1*, a classical marker of macrophages, was significantly increased in infected osteoclasts compared to not infected cells suggesting a diversion of the progenitor differentiation towards a more macrophage like phenotype (Figure 5E). Altogether, these results suggested that *C. burnetii* infection altered the expression of genes that drive the differentiation of progenitor cells into osteoclasts. In line with the phenotypic observation, this effect was dependent on the expression of a functional T4BSS by *C. burnetii*.

**Figure 5.**
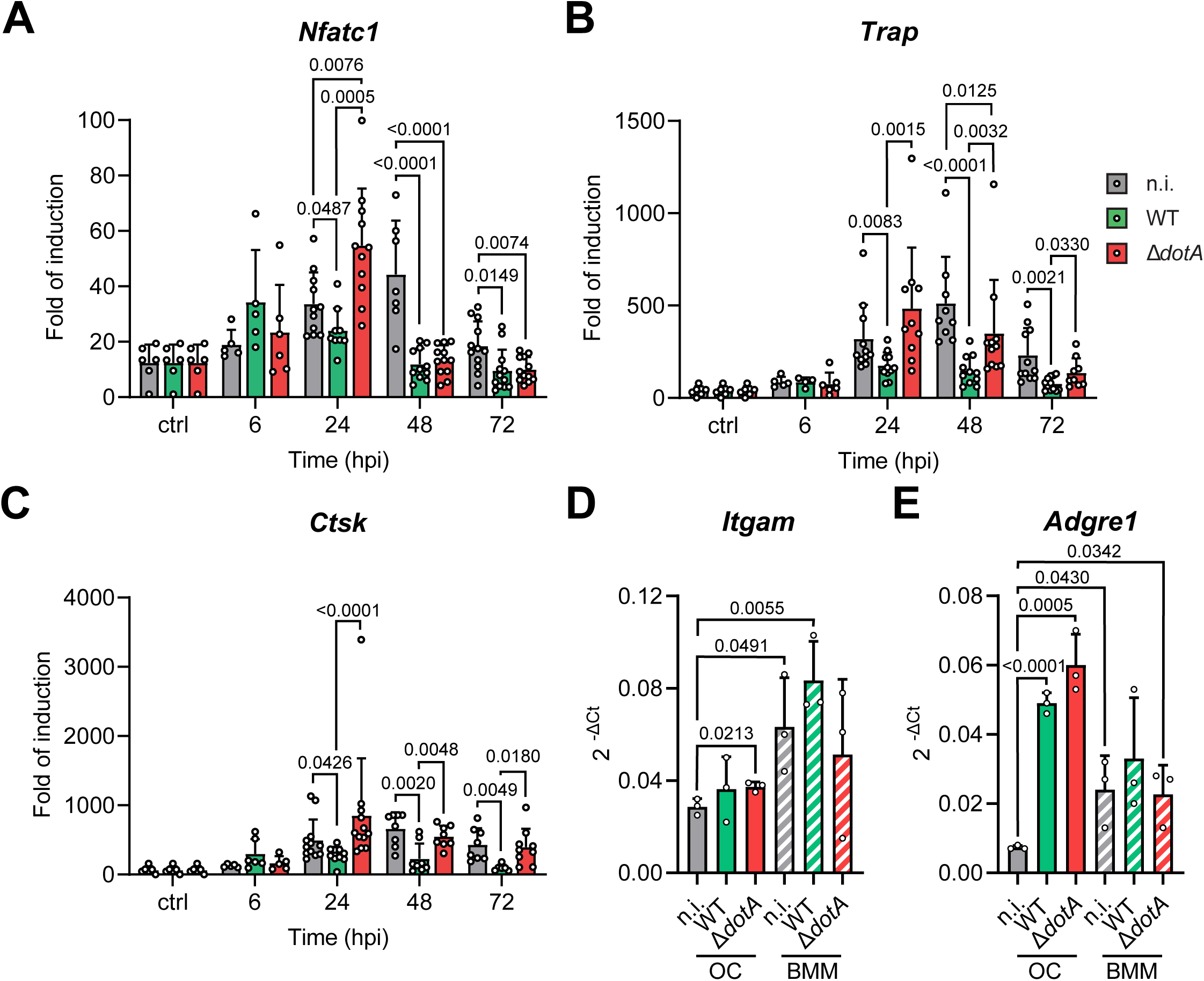
***C. burnetii* infection alters the expression of osteoclast differentiation marker genes in a DotA dependent manner. (A-C)** Osteoclasts differentiated from bone marrow progenitor cells (Fig. 3A) were infected after 2 days of differentiation with *C. burnetii* WT or Δ*dotA* at MOI 10. Uninfected osteoclasts sampled 24 h after the infection of the other cells (n.i.) were used as control. mRNA expression of osteoclast marker genes **(A)** *Nfatc1,* **(B)** *Trap* and **(C)** *Ctsk* was measured by qPCR during the differentiation in cells after 6, 24, 48 and 72 hpi (n = 6-8). **(D-E)** BM macrophages were infected with *C. burnetii* WT or Δ*dotA* at MOI 10 for 24 h. mRNA expression of macrophage marker genes **(D)** *Itgam* and **(E)** *Adgre1* was measured by qPCR in osteoclast and macrophage samples infected for 24 hpi. Uninfected osteoclasts and macrophages sampled 24 h after the infection of the other cells (n.i.) were used as control (n=3). Data are shown as mean ± SD. Circle represent individual samples.

### 4.5 Osteoclasts infected with *C. burnetii* are protected from apoptosis *in vitro*

We hypothesized that the decreased number of osteoclasts observed after infection with *C. burnetii* infection (Figure 3B) was due to a defective differentiation process (Figure 5). However, it could be also possible, that the infection induces cell death. To determine the underlying reason, the total number of cells was quantified as a proxy for cell viability (Figure 6A). In parallel, the total number of nuclei was quantified to account for the fusion events occurring during osteoclastogenesis (Figure 6B). Non-infected cells showed a decrease of the total number of cells between 24 and 72 h post-infection (Figure 6A), while keeping similar number of nuclei over this period of time (Figure 6B). In addition, the number of osteoclasts was relatively stable (Figure 3A) with increasing numbers of nuclei. This suggests that fusion events took place between osteoclasts during this period (Figure 3D). In cells infected with WT *C. burnetii*, the picture was different. More cells and more nuclei were counted over the infection time when compared to non-infected cells (Figure 6A and B). This might be due to the smaller number of osteoclasts counted in this condition (Figure 3B). Furthermore, it suggests that infected cells survived, but did not fuse with each other keeping the number of individual cells high. On the other side, infection with *C. burnetii* Δ*dotA* had a very different outcome characterized by a total number of cells similar to the non-infected cells (Figure 6A). In addition, the number of nuclei counted for *C. burneti*i Δ*dotA* infected cells was smaller than for the two other conditions (Figure 6B), pointing to a possible loss of cells. Collectively, these data suggested that infection with *C. burnetii* did not induce cell death and might even protect against it.

**Figure 6.**
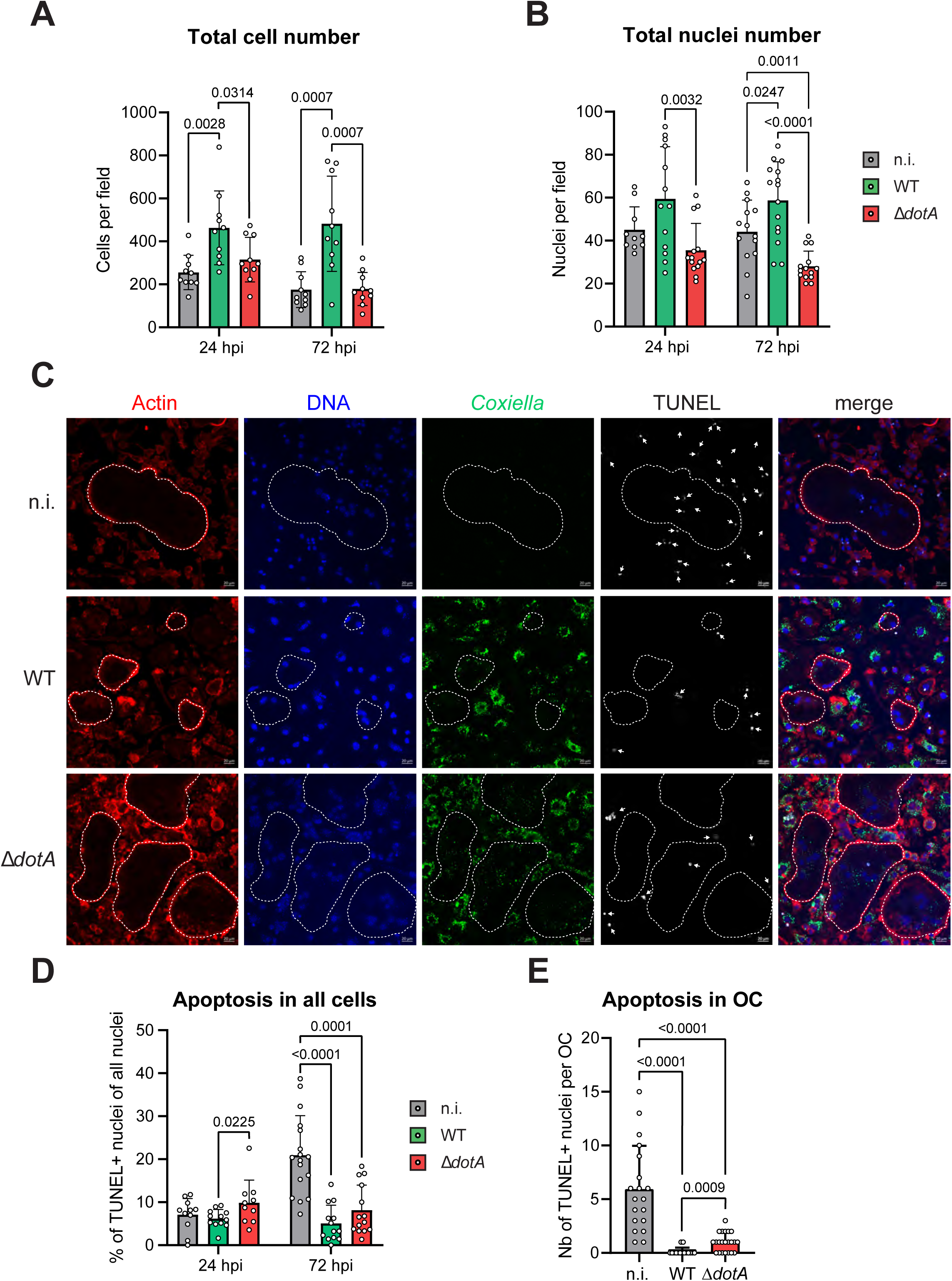
***C. burnetii* infection of osteoclasts increases their viability in a DotA dependent manner. (A)** Osteoclasts differentiated from bone marrow progenitor cells (Fig. 1A) were infected after 2 days of differentiation with *C. burnetii* WT or Δ*dotA* at MOI 10. Uninfected osteoclasts (n.i.) were used as control. Cells were stained for TRAP and analyzed by light microscopy. Total cell number was quantified with ImageJ software. **(B)** Osteoclasts previously seeded on glass coverslips were infected with *C. burnetii* WT or Δ*dotA* at MOI 10 on day 2 of differentiation. After 24 or 72 hpi, cells were stained for *Coxiella* (green - Alexa488), actin (red – Alexa647) and DNA (blue - DAPI) and imaged by fluorescent CLSFM (n=3). The total number of nuclei was quantified with ImageJ software. **(C)** Similar samples were stained in addition for *Coxiella* (green - Alexa488), actin (red – Alexa647), DNA (blue - DAPI) and TUNEL (white) and imaged by fluorescent CLSFM. Total number of nuclei and TUNEL+ nuclei were quantified with ImageJ software. Images represent the 72 hpi samples. The scale bar represents 20 µm. (n=3) **(D)** The percentage of TUNEL+ nuclei among the total number of nuclei was quantified after 24 and 72 hpi. **(E)** The number of TUNEL+ nuclei per osteoclast was quantified at 72 hpi. Data are shown as mean ± SD. Circles represent individual microscopical fields of view.

To test this possibility, we quantified the number of apoptotic cells during *in vitro* infection by TUNEL staining (Figure 6C). Less than 10% of cells were apoptotic in all conditions at 24 h post-infection. The non-infected culture harbored 21% of apoptotic nuclei among the whole cell population at 72 h post-infection. This increased apoptotic rate was not observed in cells infected with *C. burnetii* WT or Δ*dotA* (Figure 6D). Focusing on osteoclasts, we observed a similar pattern. At 72 h post-infection, non-infected osteoclasts harbored 6 apoptotic nuclei per cell (Figure 6E), which corresponds to almost half of their total nuclei (Figure 3D). This proportion drastically decreased when osteoclasts were infected with WT *C. burnetii* as no apoptotic nucleus could be observed for most cells (Figure 6E). Interestingly, apoptotic nuclei could be detected in osteoclasts infected with *C. burnetii* Δ*dotA* (Figure 6E). However, these events were still rare (1 nucleus per OC) compared to non-infected cells and represented only a minute proportion of the 30 nuclei that these cells carry when infected with the mutant strain for 3 days (Figure 3D). Noteworthy, quantification of the TUNEL staining for osteoclasts was not possible at 24 h post-infection due to the very low number of positive events at this time. Nonetheless, this experiment confirmed that infection with *C. burnetii* tend to protect cells and particularly osteoclasts against apoptosis. Even though this protection appeared to be mainly T4BSS independent, we hypothesized that among the pool of effector proteins secreted by *C. burnetii*, some might play a role in the phenotype observed during osteoclast infection.

### 4.6 Anti-apoptotic effector proteins of *C. burnetii* differently affect osteoclast differentiation and survival

Among the arsenal of effector proteins secreted by *C. burnetii*, AnkG (27, 47, 55), CaeA (25) and CaeB (24, 26) have been characterized as anti-apoptotic effector proteins. *C. burnetii* mutants lacking either AnkG or CaeB have impaired anti-apoptotic activity in HeLa and / or THP-1 cells (26, 27). Therefore, mutant strains for these two genes, WT bacteria, the Δ*dotA* mutant and heat killed bacteria were used to infect differentiating osteoclasts and the influence on OC numbers, differentiation and viability was analyzed. Heat killed bacteria were used to assess the possibility that PAMPs alone were responsible for the previously observed phenotypes. Surprisingly, the infection with the *C. burnetii* strains deficient for the two anti-apoptotic effector proteins resulted in opposite outcome. Whereas the Δ*ankG* strain failed to inhibit osteoclastogenesis (119 OC/field), the Δ*caeB* abrogated the differentiation process (8 OC/field) exceeding the WT strain inhibition (74 OC/field) (Figure 7A). In line with this result, a similar pattern was observed for the cell diameter (Figure 7B) and the number of nuclei (Figure 7C) after infection with Δ*ankG* strain producing larger osteoclast with more nuclei whereas infection with the Δ*caeB* strain reduced their size and their nuclei number. These results suggest that AnkG might inhibit osteoclastogenesis, whereas CaeB seems to enhance the differentiation process.

**Figure 7.**
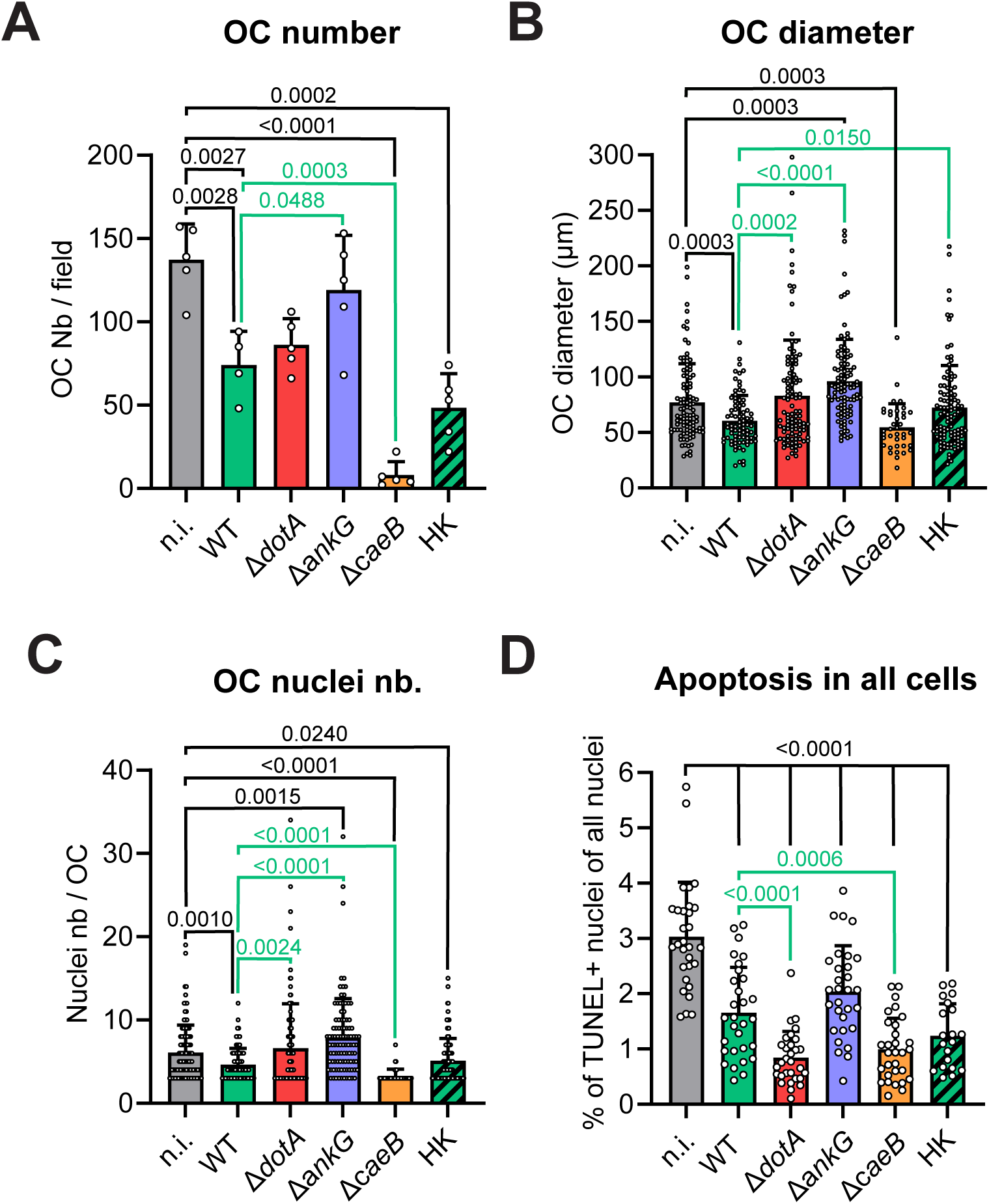
Effector proteins of *C. burnetii* affect osteoclast differentiation, but not survival. Osteoclast cultures were infected with *C. burnetii* WT, Δ*dotA*, Δ*ankG*, Δ*caeB* or heat-killed WT (HK) after 2 days of RANKL stimulation. Uninfected osteoclasts (n.i.) were used as control. Osteoclast differentiation was quantified 72 hpi by staining for TRAP. Cells were analyzed by light microscopy. **(A)** Osteoclasts (TRAP+ cells with ≥3 nuclei) were counted with imageJ software (n=3). **(B)** Similar approaches were used to measure the osteoclast diameter and **(C)** the number of nucleus per field as a proxy for cell survival. **(D)** Similar infection experiments were carried out on osteoclasts previously seeded on glass coverslip. After 72 hpi, cells were stained for *Coxiella* (green - Alexa488), actin (red – Alexa647), DNA (blue - DAPI) and TUNEL (white) and imaged by fluorescent CLSFM. Total number of nuclei and TUNEL+ nuclei were quantified with ImageJ software. The percentage of TUNEL+ nuclei among the total number of nuclei was quantified (n=2). Data are shown as mean ± SD. Circle represent individual microscopical fields of view.

The analysis of the effect of these effector proteins on the protection of the cell in culture from apoptosis showed that the absence of AnkG did not alter the protection provided by the WT strain. However, the absence of CaeB secretion had a surprising enhancing protective effect (Figure 7D). These data demonstrate that neither AnkG nor CaeB are important for *C. burnetii*-mediated anti-apoptotic activity during osteoclast infection. The anti-apoptotic activity might be explained by the results obtained with the heat-killed WT *C. burnetii*. Indeed, the phagocytosis of dead bacteria by osteoclasts induced a protection from apoptosis like the one observed with WT *C. burnetii* (Figure 7D).

### 4.7 Infection with *C. burnetii* inhibits the bone-resorbing function of osteoclasts *in vitro*

Considering the alteration of the differentiation of osteoclasts following infection with *C. burnetii*, we hypothesized that the bone resorption capacity of infected cells could also be impaired. Therefore, we tested their functional activity in an *in vitro* resorption assay. There, osteoclasts are seeded and differentiated in well plates coated with a layer of calcium phosphate as substrate. Differentiating osteoclasts were infected with *C. burnetii* strains following a protocol similar to the previous experiments (Figure 2A). The quantity of resorbed calcium phosphate was measured by light microscopy at the end of the experiment. At 5 days post-infection, 21% of the surface of the well was resorbed by the non-infected cells (Figure 8). This resorption activity was reduced when osteoclasts were infected with WT *C. burnetii* to 4% of the resorbed surface. However, osteoclasts infected with *C. burnetii* Δ*dotA* developed a strong resorption activity, which was even greater than that of the non-infected cells. This result supported the correlation between the osteoclast number, their size and their resorption activity. While cells infected with WT *C. burnetii* were not properly differentiating and harbored a weak resorption activity, the cells infected with the Δ*dotA* mutant grew larger and developed a stronger lytic activity (Figure 3B, 3D and 8).

**Figure 8.**
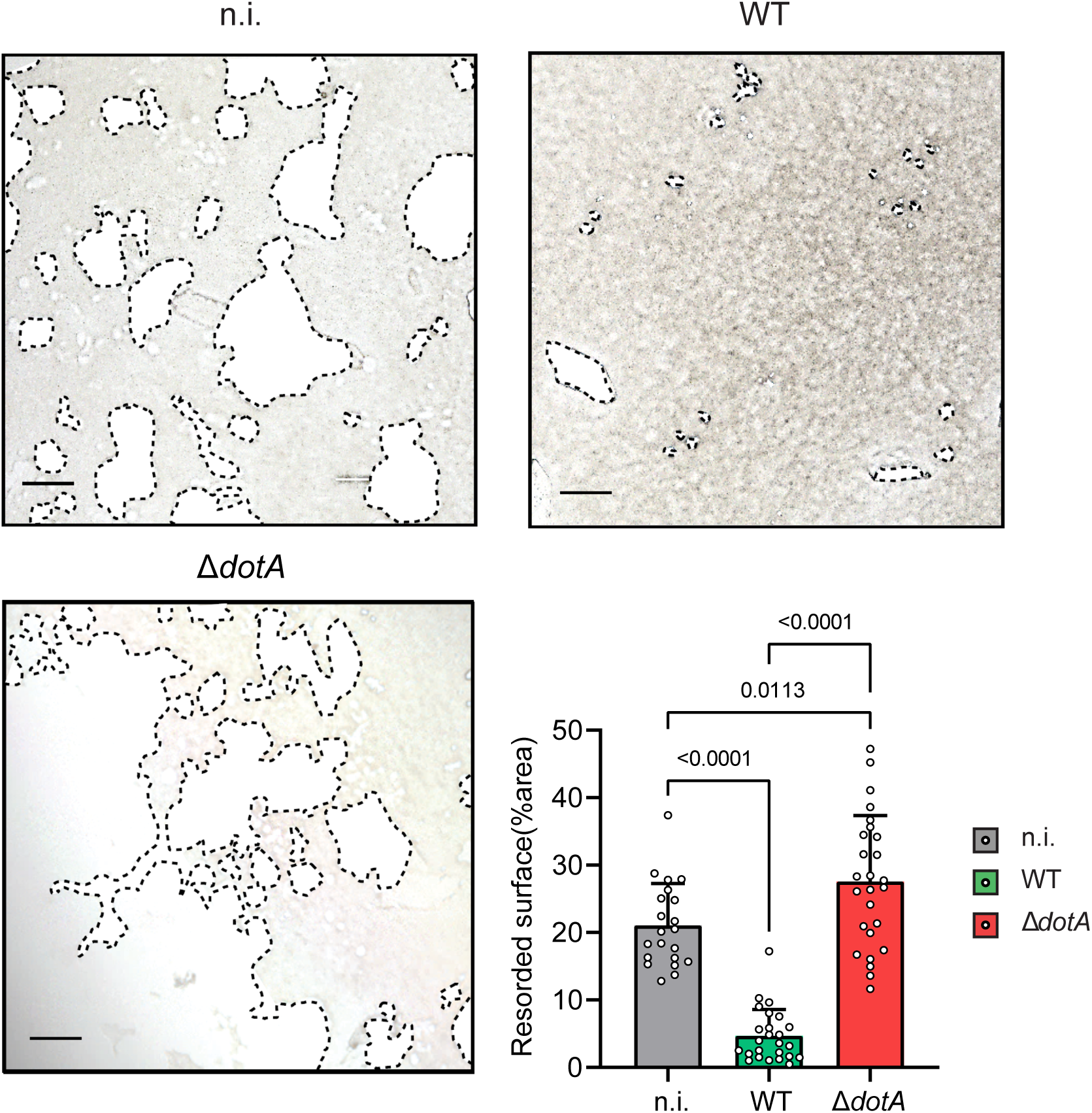
***C. burnetii* infection of osteoclasts inhibits their bone resorption activity in a DotA dependent manner.** The degradation activity of osteoclasts was measured with cells seeded on a collagen embedded plate using a differentiation and infection protocol similar to figure 1A. Infection was realized on day 2 of differentiation. Degradation activity was analyzed on day 7 (5 dpi). Plates were imaged by light microscopy. Area of collagen degradation (resorption pit) was quantified with ImageJ (3 mice for each condition; 5 wells per mice and condition). Scale bar represents 100 μm. Data are shown as mean ± SD. Circle represent individual microscopical fields of view.

### 4.8 The inflammatory response of osteoclasts is repressed by the T4SS of *C. burnetii*

To continue our developmental and functional characterization of infected osteoclasts, we investigated the immunological response of these cells to infection. At first, the expression of pro-inflammatory cytokine genes, such as *Tnf*, were analyzed. For all time points covering up to 72 h post-infection, we observed a limited induction of *Tnf* with no substantial difference between the cells infected with WT *C. burnetii* and the non-infected cells (Figure 9A). However, cells infected with *C. burnetii* Δ*dotA* expressed early on significantly more *Tnf* than cells infected with WT bacteria or non-infected cells. The limited gene expression did not translate into a measurable amount of secreted TNF (Figure 9B). Regarding *Il1b*, only the cells infected with *C. burnetii* Δ*dotA* induced a limited gene expression (Figure 9C) that translated in a minute amount of secreted cytokine for this particular condition (Figure 9D). While *Tnf* and *Il1b* showed generally low level of induction, Il6 was strongly induced by infection with *C. burnetii*. Osteoclasts infected with the WT bacteria had a quick and sustained induction of *Il6*, which peaked at 48 h post-infection and rescinded completely at 72 h post-infection (Figure 9E). Infection with the Δ*dotA* mutant induced an earlier wave of *Il6* expression than for the cells infected with the WT *C. burnetii*. Nonetheless, the levels of induction were in the same range for both bacterial strains. This was confirmed by the measurement of a similar level of secreted IL-6 in both conditions (Figure 9F). Finally, we measured the level of expression of the anti-inflammatory cytokine gene *Il10*. The expression pattern of *Il10* was similar to the analyzed pro-inflammatory cytokines, as it was slightly higher induced by the infection with WT *C. burnetii* in comparison to non-infected cells during the course of infection (Figure 9G). However, infection with *C. burnetii* Δ*dotA* induced a higher level of *Il10* expression than infection with the WT bacteria at early time point. This kinetic difference appeared to even out when we analyzed the quantity of IL-10 secreted after 24h (Figure 9H). Altogether, our data suggested that *in vitro* osteoclast cultures have a rather limited inflammatory response to infection with WT *C. burnetii*. In addition, these results suggested that infection with the Δ*dotA* mutant might induce a stronger pro-inflammatory response than the infection with the WT bacteria.

**Figure 9.**
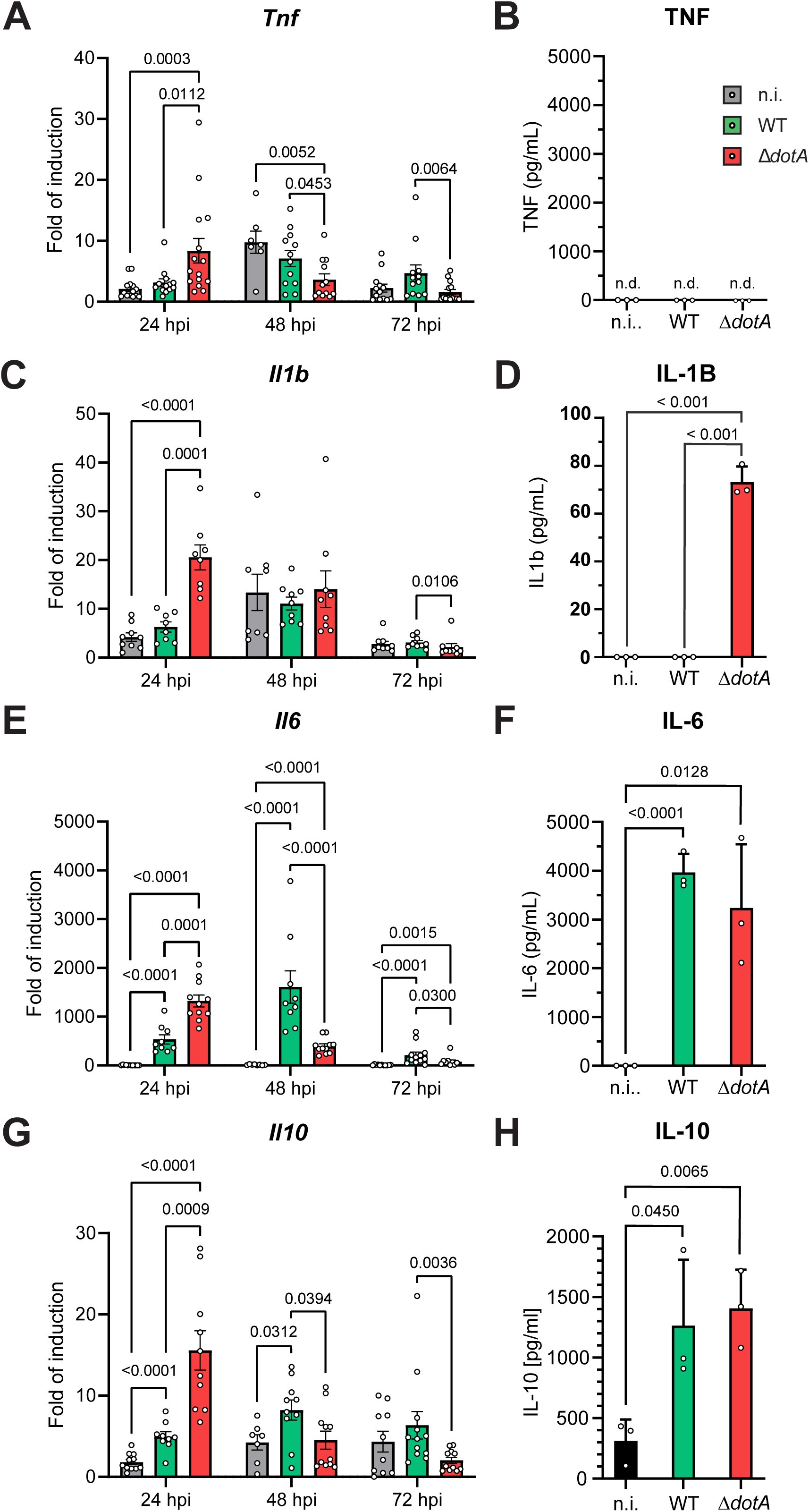
***C. burnetii* needs DotA to restrain the cytokine production of osteoclasts after infection.** Osteoclasts differentiated from bone marrow progenitor cells (Fig. 1A) were infected after 2 days of differentiation with *C. burnetii* WT or Δ*dotA* at MOI 10. Uninfected osteoclasts sampled 24 h after the infection of the other cells (n.i.) were used as control. mRNA expression of cytokine genes **(A)** *TNF,* **(C)** *Il1b*, **(E)** *Il6* and **(G)** *Il10* was measured by qPCR during the differentiation in cells after 24, 48 and 72 hpi (n=6). Secretion of cytokine **(B)** TNF, **(D)** IL-1B, **(F)** IL-6 and **(H)** IL-10 was also quantified in the supernatant of the cells 24 hpi (n=3). Data are shown as mean ± SD. Circle represent individual samples.

## 5 Discussion

The mechanisms of persistence of *C. burnetii* within humans during chronic Q fever remain elusive. Long-term studies of cohorts of patients suffering chronic Q fever suggested that the persistence was largely linked to the presence of *C. burnetii* in the bone marrow (33). The findings presented in this report identify osteoclasts as candidate to perpetuate the infection in this organ.

First, we demonstrated that osteoclasts, as phagocytic cells, were efficiently infected with *C. burnetii* (Figure 2). This was expected knowing that the two integrins (11, 56) used by other phagocytes to capture *C. burnetii* are also present at the surface of osteoclasts. In addition to its phagocytic role, integrin α5β3 (CD51/CD61) is indeed highly expressed by osteoclasts to sense their extracellular matrix and to provide, in response, a pro-survival stimulus (57). Similarly, osteoclasts express the integrin αMβ2 (CD11b/CD18) like all other myeloid cells. The presence of this phagocytic complement receptor is required for completing osteoclastogenesis (58). While the independent role of each host integrin was not addressed in this study, a role of the T4BSS could be excluded in the phagocytic process. This result was in line with the expression kinetic of the secretion system described in other cells within which it becomes functional only after the maturation of the phagolysosome, around 8-24 h post infection (19, 59). Importantly, we could establish a role for the T4BSS in manipulating the host cell and allowing the intracellular growth of the bacteria in osteoclasts (20). While the burden of WT *C. burnetii* increased over time like in other myeloid cells (60), the Δ*dotA* mutant could not grow in osteoclasts (Figure 2).

Although the T4BBS takes up to 8 to 24 hpi to be functional (19), it is critical for the bacteria to inject its arsenal of virulence factors. This includes anti-apoptotic virulence factors that maintain host cell survival by counteracting the stress induced by the developing intracellular infection (61). This pro-survival effect of *C. burnetii* infection was also observed in osteoclasts as illustrated by the strongly decreased number of apoptotic nucleus observed in the later phase of the infection (Figure 6). Surprisingly, the T4BSS did not have a preponderant role in this process as the Δ*dotA* mutant protected cells at level similar to the WT C*. burnetii*. This result diverged from the results obtained in macrophages and other non-myeloid cells for which the injection of the virulence factors is required for the protection (62) (20). Indeed, our data demonstrate that the anti-apoptotic effector proteins AnkG and CaeB are dispensable for *C. burnetii*-mediated inhibition of cell death of osteoclasts (Figure 7D). Importantly, in case of osteoclasts, the phagocytosis of live and heat-killed *C. burnetii* alone had an anti-apoptotic effect similar to the one observed in the context of other bacterial infections such as with *Staphylococcus aureus* (63). While AnkG and CaeB are not involved in inhibition of apoptosis in osteoclasts, they might affect osteoclast development, which need further investigations. Furthermore, heat-killed *C. burnetii* alters the differentiation process (Figure 7A and B). Indeed, pathogen-and danger-associated molecular patterns (PAMPs and DAMPs) can enhance the differentiation and survival of osteoclasts either by directly skewing their metabolism toward oxidative phosphorylation (64) or via paracrine and autocrine secretion of proinflammatory cytokines like TNF and IL-6 (65). The later could partly explain the pro-survival effect observed in osteoclasts as we observed the expression of these cytokines after infection with *C. burnetii* (Figure 5). Interestingly, the induction of these pro-inflammatory genes was earlier and stronger in osteoclasts infected with the Δ*dotA* mutant in comparison to WT *C. burnetii*, despite being decorated with identical PAMPs and infecting similarly osteoclasts. This suggests that an efficient infection, that relies on the expression of T4BSS, could overtake the infected cell signaling pathways to reduce the proinflammatory response, as observed for IL-1B, giving time to the bacteria to adapt and proliferate. Secretion of IL-10 after infection might also participate in the reduction of this inflammatory response. As a consequence, the delayed increase of pro-inflammatory cytokine expression following infection with WT *C. burnetii* might be due to the increased bacterial burden promoted by the T4BSS.

Another consequence of the infection was the reduction of the osteoclast differentiation process. Osteoclasts infected with WT *C. burnetii* were smaller, contained less nuclei and expressed less differentiation markers than uninfected cells (Figure 3). This indicated a reduction of the fusion events in osteoclasts carrying replicative infection as increased cell death was previously excluded as a possible cause of this phenotype. Surprisingly, the differentiation of osteoclasts infected with the Δ*dotA* strain was not altered. The differentiation was even enhanced as a higher number of nuclei was counted in osteoclasts infected with Δ*dotA C. burnetii* than in uninfected osteoclasts. We hypothesize two reasons for this increase of fusion events: i) Either infection by the Δ*dotA* strain induced a more osteoclastogenic cytokine profile than the WT strain enhancing the osteoclastogenesis program, or ii) the anti-apoptotic effect induced by infection allowed the cells to encounter more osteoclast progenitors and subsequently more fusion events. Regarding the first hypothesis, we measured similar amount of IL-10, a cytokine known to inhibits osteoclastogenesis by reducing the expression and the nuclear translocation of NFATc1 (66, 67), and IL-6, a osteoclastogenic cytokine (65). Nonetheless, we cannot exclude that an imbalance of other cytokines could explain this difference. Independently of the pathway taken, the giant osteoclasts resulting from the infection with the Δ*dotA* strain exhibited the strongest bone resorbing activity *in vitro* whereas the few ones who developed after the infection with the WT strain had a reduced activity despite the fact that both strains provided protection from apoptosis. Altogether, the *in vitro* data presented in this report demonstrated that *C. burnetii* can infect and replicate within osteoclasts. In addition, the infection altered the differentiation and function of osteoclasts in a manner depending on the expression of the T4BSS.

This suggested a putative participation of osteoclasts during Q fever. This hypothesis was validated by our murine model of Q fever. Mice are naturally resistant to *C. burnetii*. However, mice deficient for the TLR and IL1R adaptor MyD88 are susceptible to the infection. Systemic bacterial burden is indeed increased in these mice over the course of infection (49). Recapitulating this model, we observed that the bone marrow of MyD88 deficient mice also show an increase of susceptibility to the infection 5 dpi. The immune-histological analysis of bones of infected mice revealed that trabeculae-lining osteoclasts could also be infected *in vivo* by *C. burnetii* (Figure 1). Whereas the proportion of infected osteoclast was similar between immune-sufficient and-deficient mice, the increased number of infected bone marrow cells in immune-deficient mice correlated with their increased bacterial burden in this organ. Indeed, alteration of the innate immune system like in this murine model of Q fever might also favor the spreading of *C. burnetii* to the bone marrow in infected patients (53). This newly identified cellular niche could be of importance for the persistence of *C. burnetii* during chronic Q fever as bone marrow cells have been linked to the persistence of other bacterial infections like with *Mycobacterium tuberculosis* or *Staphylococcus aureus* (38, 42, 68, 69). In addition, this could have new implications for chronic Q fever. As infection reduces the bone resorbing activity of osteoclasts and fully blocks the differentiation of monocytic precursor into osteoclast, the bone structure could be altered leading to an osteopetrotic phenotype in chronic Q fever patients. However, there are some limitations of our study, including the infection model. Intraperitoneal infection used in this study might not mimic the natural route of infection. Furthermore, the timing of our bone sampling might not represent the persistent stage of infection. Future study of the persistent phase of intratracheal-infection will help us to decipher the role of osteoclasts in this process. Altogether, this report opens new avenues for investigating therapeutic interventions targeting *C. burnetii* persisting in osteoclasts to treat chronic Q fever.

## 7 Author Contributions

D.S. and A.L. conceived and coordinated the study. C.L. designed, performed and analyzed the *in vitro* experiments. J. S.-L. performed *in vitro* experiments. F.A. analyzed *in vitro* and *in vivo* experiments. X.S. performed and analyzed immunohistofluorescence of in vivo samples. R.L. conceived and coordinated the *in vivo* experiments. M.N.A.A.S. designed and performed the *in vivo* experiments, which were analyzed by C.L. with the help of Y.J. and A.B.. D.S. wrote the manuscript with the support of A.L.. All authors reviewed and approved the final manuscript.

## 9 Acknowledgments

We would like to thanks Claudia Feulner for her constant technical support and Barbara Bodendorfer for help with mouse infection.

